# Highly sensitive reverse transcription loop-mediated isothermal amplification for detecting and identifying groundnut ringspot and tomato spotted wilt orthotospoviruses

**DOI:** 10.1101/2024.11.29.625922

**Authors:** Soledad de Breuil, Salvador López, Carolina Dottori, Claudia Nome, Scott Adkins

**Affiliations:** Instituto Nacional de Tecnología Agropecuaria, Centro de Investigaciones Agropecuarias, Instituto de Patología Vegetal Ing. Agr. Sergio Fernando Nome (INTA-CIAP-IPAVE), Av. 11 de Septiembre 4755, X5014MGO, Córdoba, Argentina; Consejo Nacional de Investigaciones Científicas y Técnicas (CONICET), Unidad de Fitopatología y Modelización Agrícola (UFYMA), Av. 11 de Septiembre 4755, X5014MGO, Córdoba, Argentina; United States Department of Agriculture, Agricultural Research Service (USDA-ARS), U.S. Horticultural Research Laboratory, Fort Pierce, FL 3494, United States

**Author notes:** Corresponding author: Soledad de Breuil.

**Keywords:** Molecular identification, RT-LAMP, *Orthotospovirus*, tomato, pepper, peanut

## Abstract

*Orthotospovirus arachianuli* (groundnut ringspot virus, GRSV) and *Orthotospovirus tomatomaculae* (tomato spotted wilt virus, TSWV) represent a threat to several agricultural crops worldwide. The management of the diseases they cause is particularly challenging due to their wide host range and transmission by thrips vectors. Accurate diagnosis and early detection are crucial, especially considering the limitations of conventional serological and molecular methods. This study introduces reverse transcription loop-mediated isothermal amplification (RT-LAMP) as a reliable alternative for detecting and identifying GRSV and TSWV in tomato, pepper, and peanut plants. The designed primer sets specifically targeted and amplified fragments of the viral RNA S segment. RT-LAMP reactions produced negative results in the healthy plants and exhibited no cross-reactivity with closely related orthotospoviruses or viruses from other genera. Compared to RT-PCR, RT-LAMP exhibited superior sensitivity, detecting viral RNA at concentrations 100 times lower. Field testing on symptomatic plants further confirmed the effectiveness of this technique. Overall, this study highlights RT-LAMP as a rapid, species-specific, and highly sensitive tool for the early detection of GRSV and TSWV in economically important crops, offering a promising solution for improved disease management.

---

Viral plant diseases are among the leading causes of severe epidemics reported worldwide (Patil, 2021). Among plant viruses, the genus *Orthotospovirus* (family *Tospoviridae*) comprises species considered highly harmful to agricultural production. *Orthotospovirus arachianuli* (groundnut ringspot virus, GRSV) is an emerging orthotospovirus that threatens the quality and yield of several important crops, including vegetables and ornamentals in Argentina, Brazil, China, Ghana, Kenya, Paraguay, South Africa, and the United States (EPPO 2024). In the United States, GRSV has been found only as an interspecies recombinant strain between GRSV and *Orthotospovirus tomatoflavi* (tomato chlorotic spot virus, TCSV), designed L_G_M_T_S_G_ (Webster et al., 2011).

*Orthotospovirus tomatomaculae* (tomato spotted wilt virus, TSWV), the type species of the genus, is the most widespread orthothospovirus and infects over 1,000 species of cultivated and wild plants worldwide. Due to the extensive damage caused by TSWV, it has been classified as one of the most devastating viral pathogens affecting agricultural production (Ranawaka et al., 2020).

Effectively managing orthotospoviruses poses significant challenges due to their wide host range and persistent, propagative transmission by thrips vectors. Addressing diseases caused by orthotospoviruses necessitates the implementation of integrated management strategies, including cultural practices (such as optimizing sowing dates, adjusting planting density, and removing infected plants), pest control measures, and the use of virus-resistant germplasm, together with strategies aimed at modulating thrips feeding behavior (Oliver and Whitfield, 2016). In this context, effective prevention and disease management rely heavily on the accurate identification of the etiological agent and the early detection of affected plants (Roossinck 2013). Viral diseases are typically identified using one or more methods, including 1) plant symptoms, 2) electron microscopy, 3) detection of target proteins, or 4) identification of target nucleic acids. The symptoms induced by GRSV, TCSV, and TSWV in infected plants are often indistinguishable, and serological tests frequently exhibit cross-reactions due to the high degree of homology in the amino acid sequences of their nucleoproteins (de Avila et al., 1990). Therefore, molecular techniques targeting the viral genome are essential for the precise identification of viral species.

The genome of orthotospoviruses consists of three RNA segments, designated as L (large), M (medium), and S (small). The L RNA is of negative polarity and encodes the RNA-dependent RNA polymerase (RdRp). The M and S RNA segments have an ambisense configuration, each containing two non-overlapping open reading frames (ORFs). The M RNA encodes the precursor of two glycoproteins (Gn and Gc) and a non-structural protein (NSm) essential for cell-to-cell movement in plants. The S RNA encodes the nucleoprotein (N) and the non-structural protein (NSs), which suppresses gene silencing (King et al., 2012).

While reverse transcription-polymerase chain reaction (RT-PCR) is widely recognized as a highly effective method for detecting RNA plant viruses, it has inherent limitations. These include the requirement for expensive equipment, extended processing times, and the need for highly trained personnel. Additionally, polymerase activity can be inhibited by compounds such as polyphenols and polysaccharides, which are often present in plant tissues, potentially leading to false-negative results. Another significant drawback of PCR is its high cost, which arises from materials required for quality control, nucleic acid purification, contamination prevention during test preparation, and result interpretation (Le and Vu, 2017). Loop-mediated isothermal amplification (LAMP) has emerged as a highly efficient, specific, and cost-effective alternative amplification technique that addresses many of the drawbacks associated with PCR (Notomi et al., 2000). In particular, reverse transcription LAMP (RT-LAMP) has been successfully applied for the detection of RNA viruses (Panno et al., 2020). LAMP does not require specialized equipment or highly trained personnel, yet it offers high specificity by utilizing a set of primers designed to recognize independent fragments of a target region. Typically, the primer set consists of six primers: four primary primers that target specific sections of the nucleic acid—forward and backward outer primers (F3 and B3), and forward and backward inner primers (FIP and BIP)—along with two loop primers (LF and LB) that enhance the reaction speed, specificity, and sensitivity. The amplification process is facilitated by the *Bst* DNA polymerase, a chain-displacement enzyme derived from *Geobacillus stearothermophilus*. This enzyme promotes the formation of loop structures by internal primers, enabling rapid, self-priming amplification under isothermal conditions. Additionally, the robustness of *Bst* polymerase reduces the impact of inhibitors commonly present in plant tissue extracts (Nagamine et al., 2002).

The objective of this study was to develop and implement an RT-LAMP method for the simple, rapid, and accurate detection of GRSV and TSWV in infected plants. The work focused on tomato, pepper, and peanut crops, which are globally important and frequently impacted by these viral pathogens.

The complete nucleotide sequences of the N gene for GRSV, TCSV, and TSWV were retrieved from GenBank (National Center for Biotechnology Information). These sequences were aligned using MUSCLE and consensus sequences were generated for each viral species. The resulting consensus sequences were mapped to each other, and a nucleotide homology analysis was performed using Geneious Prime (version 2022.0.2). The consensus sequences for GRSV and TSWV were specifically utilized to design LAMP primers. Primer sets (FIP, BIP, F3, B3, LF, and LB) were designed using the NEB LAMP Primer Design Tool (https://lamp.neb.com/), set for AT-rich sequences. The selected primers are listed in Table 1, and their corresponding target genomic regions are shown in Fig. 1.

**Fig 1.**
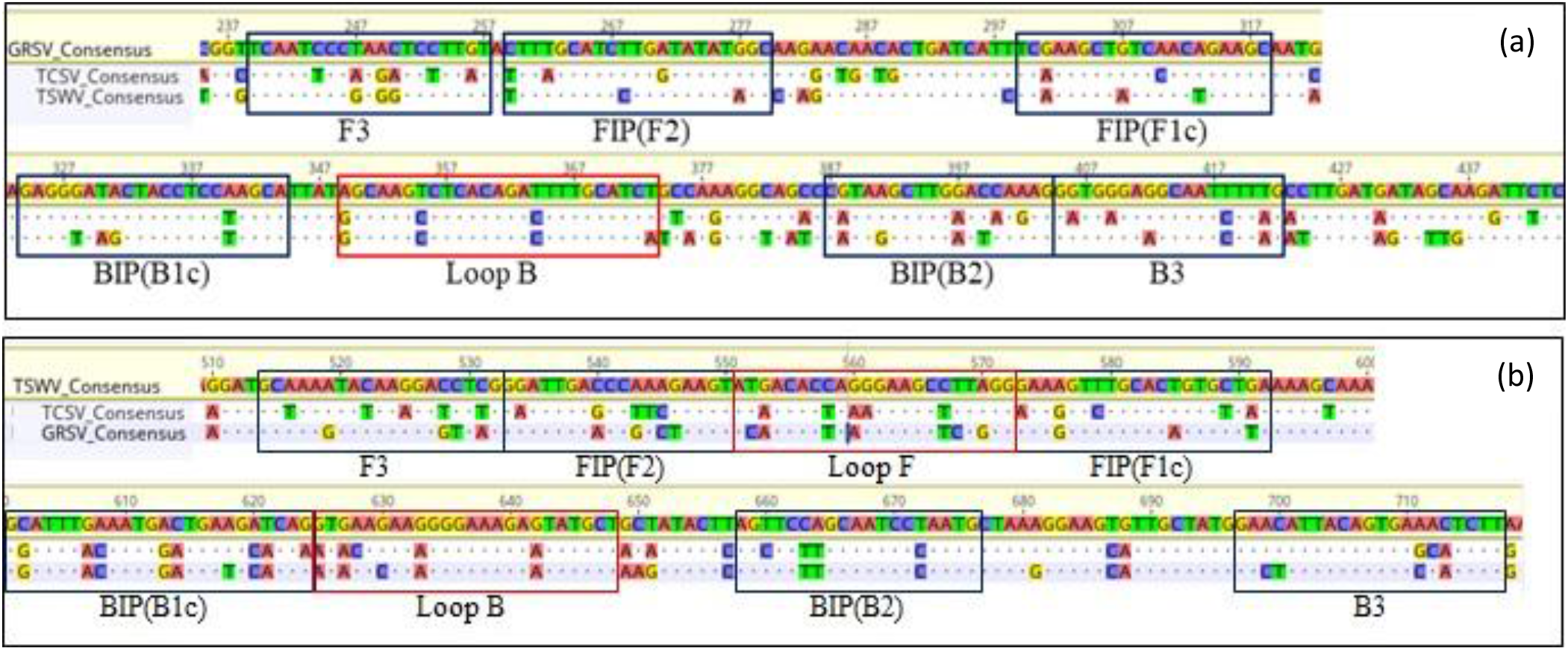
Mapping of genome consensus sequences of the nucleoprotein (N) gene from groundnut ringspot virus (GRSV), tomato chlorotic spot virus (TCSV), and tomato spotted wilt virus (TSWV). (a) Sequence regions targeted by the primer set for amplifying GRSV. (b) Sequence regions targeted by the primer set for amplifying TSWV.

**Table 1.**
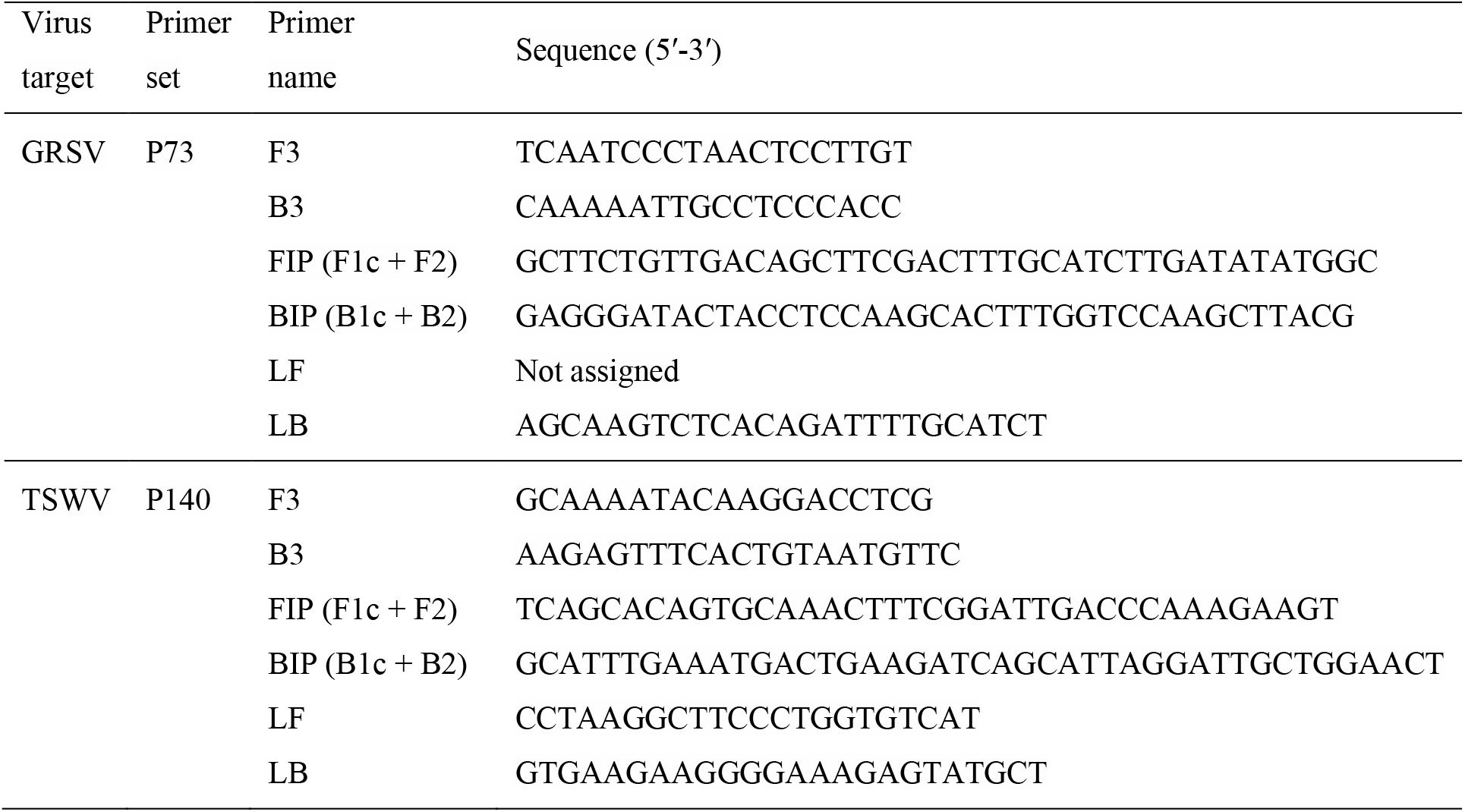
Primer sets and sequences of RT-LAMP primers for detecting groundnut ringspot virus (GRSV) and tomato spotted wilt virus (TSWV).

The sources of inocula included GRSV (L_G_M_T_S_G_ isolate), TCSV, and TSWV, collected from tomatoes in Florida, USA. These were maintained in tomato, pepper, and *Datura stramonium* via mechanical inoculations, according to Webster et al. (2011). The genome of the GRSV-L_G_M_T_S_G_ isolate comprises RNA L and RNA S from GRSV, and RNA M from TCSV (Webster et al., 2011). Virus-infected and healthy plants served as positive and negative controls, respectively, in all assays.

The presence of single viral infections and the health status of control plants were confirmed by RT-PCR. Total RNA was extracted from plant tissues using the RNeasy Plant Mini Kit (Qiagen Inc.), following the manufacturer’s protocol. RNA quality and yield were assessed via spectrophotometry, and samples were stored at -20°C until use. First-strand cDNA was synthesized from 500 ng/µl of total RNA using random hexamers and LunaScript RT Master Mix Kit (Primer Free) (New England Biolabs Inc.) at 55°C for 10 min. The resulting cDNA (0.5 µl per reaction) was subjected to PCR using GoTaq Green Master Mix (Promega Corp.) with specific primers for GRSV (GRSV-N-v/GRSV-N-vc), TCSV (TCSV-3’Lv/TCSV-3’Lvc), and TSWV (TSWV722/TSWV723) (Adkins and Rosskopf, 2002; Webster et al., 2011, 2013). The cycling parameters used in the PCR reactions were those described by Tantiwanich et al. (2018). RT efficiency was verified by amplifying a 181 bp fragment of the conserved mitochondrial nad5 gene using nad5-s/nad5-as primers (Menzel et al., 2002). PCR products were assessed by electrophoresis in 1.2% agarose gels stained with SYBR Safe DNA Gel Stain (Invitrogen). RT-PCR confirmed the presence of single viral infections in inoculated plants and the absence of viruses in healthy controls.

RT-LAMP reactions were conducted using WarmStart Colorimetric LAMP 2X Master Mix with UDG (New England Biolabs Inc.). Each reaction (20 μl) included 10 μl of LAMP 2X master mix, 2 μl of 10X primer mix (2 μM F3 and B3, 16 μM FIP and BIP, and 4 μM LF and LB), 1 μl of plant RNA, and 7 μl of nuclease-free water. Samples were incubated at 65°C for 30 min, and results were visually interpreted. A yellow color indicated the presence of viral RNA, whereas pink indicated a negative result. Each RT-LAMP assay was repeated three times.

The RT-LAMP assays, using primers designed in this study (P-73 for GRSV and P-140 for TSWV), successfully detected GRSV and TSWV in RNA extracted from infected tomato, pepper, peanut, and Datura stramonium plants. Primer specificity was confirmed by testing total RNA from control plants infected individually with GRSV, TSWV, and TCSV. No cross-reactivity was observed (Fig. 2). Additionally, primers were tested against RNA from healthy tomato, pepper, and peanut plants, and plants infected with impatiens necrotic spot virus (INSV, genus *Orthotospovirus*), cucumber mosaic virus (CMV, genus *Cucumovirus*), tobacco mosaic virus (TMV, genus *Tobamovirus*), papaya ringspot virus (PRSV, genus *Potyvirus*), colombian datura virus (CDV, genus *Potyvirus*) and tomato yellow leaf curl virus (TYLCV, genus *Begomovirus*). RT-LAMP showed no false positives, demonstrating the high specificity of the assays for GRSV and TSWV.

**Fig 2.**
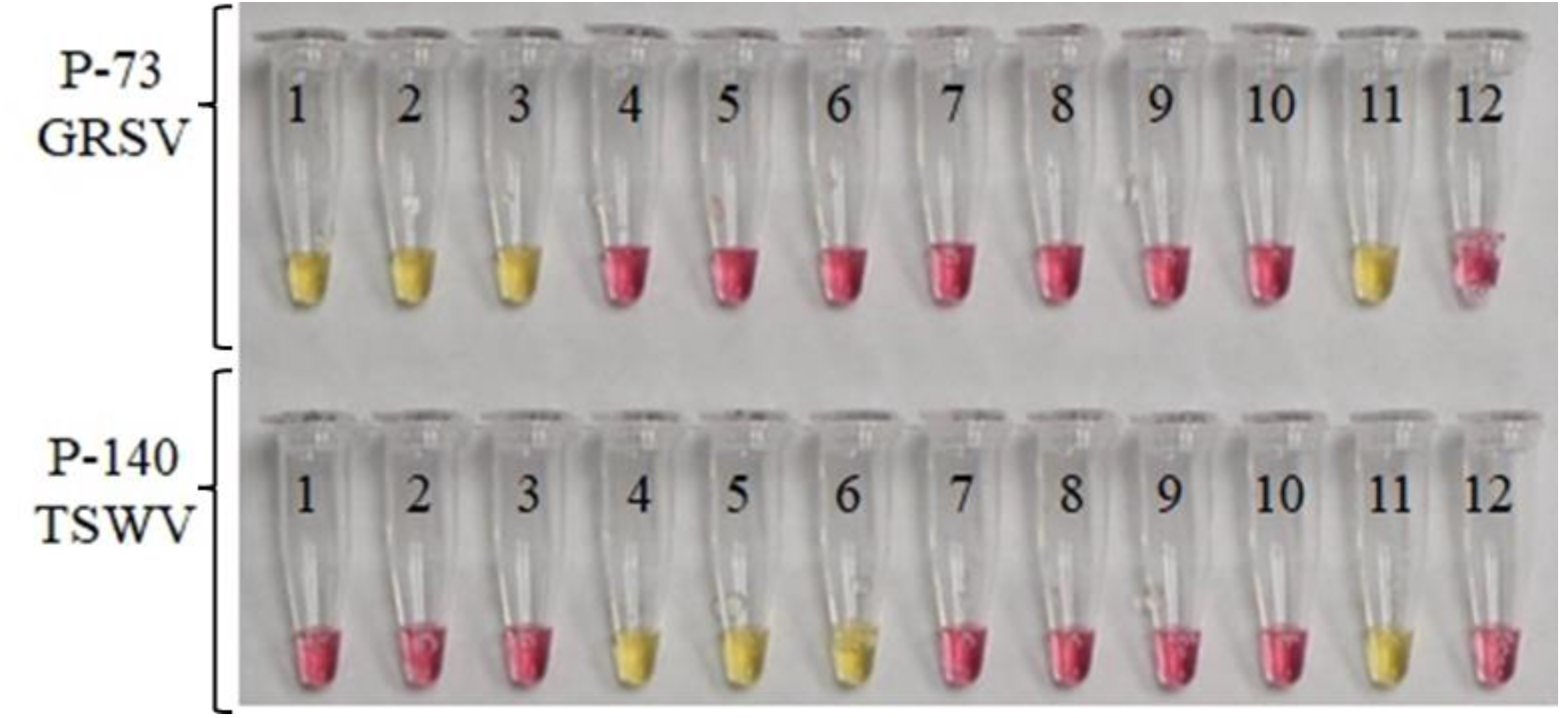
Specificity of P-73 and P-140 primer sets in RT-LAMP for detecting groundnut ringspot virus (GRSV) and tomato spotted wilt virus (TSWV), respectively. Tubes 1-3, RNA from plants tested positive for GRSV; tubes 4-6, RNA from plants tested positive for TSWV; tubes 7-9, RNA from plants tested positive for TCSV; tube 10: negative control (healthy plant); tube 11, positive control; tube 12, no template control (water).

The sensitivity of the RT-LAMP method was compared to conventional RT-PCR. Peanut plants diagnosed as positive for GRSV and TSWV were stored at -80°C for at least 1 hour and then processed for total RNA extraction as described above. To evaluate sensitivity, 10-fold serial dilutions were prepared, starting from an initial RNA concentration of 500 ng/μl. These dilutions were subjected to RT-PCR and RT-LAMP analyses, using 1 μl of RNA as input and following the protocols previously described. The RT-LAMP products were analyzed on a 2.5% agarose gel stained with SYBR Safe DNA Gel Stain (Invitrogen, USA), under an extraction hood.

RT-LAMP demonstrated a sensitivity 100 times greater than RT-PCR for detecting both GRSV and TSWV, consistent with findings reported in other studies (Bhat et al., 2022). In RT-PCR, a band of the expected size (∼595 bp) corresponding to GRSV was detectable in reactions initiated with 0.5 ng/μl of total RNA. However, the bands became progressively weaker at lower RNA concentrations, making detection challenging (Fig. 3a). In contrast, RT-LAMP reactions, assessed through colorimetric evaluation, produced well-defined positive results at RNA concentrations as low as 5 pg/μl (Fig. 3b). Furthermore, RT-LAMP fragments for GRSV were clearly visible on agarose gel down to 5 pg/μl, aligning with the colorimetric observations (Fig. 3c). Comparable results were obtained for TSWV detection. RT-PCR generated the expected band (∼620 bp) with RNA concentrations as low as 5 ng/μl (Fig. 3d), whereas RT-LAMP produced easily distinguishable yellow reactions at RNA concentrations as low as 0.05 ng/μl (Fig. 3e). The yellow color in RT-LAMP reactions correlated with amplified fragments observed on agarose gel (Fig. 3f).

**Fig 3.**
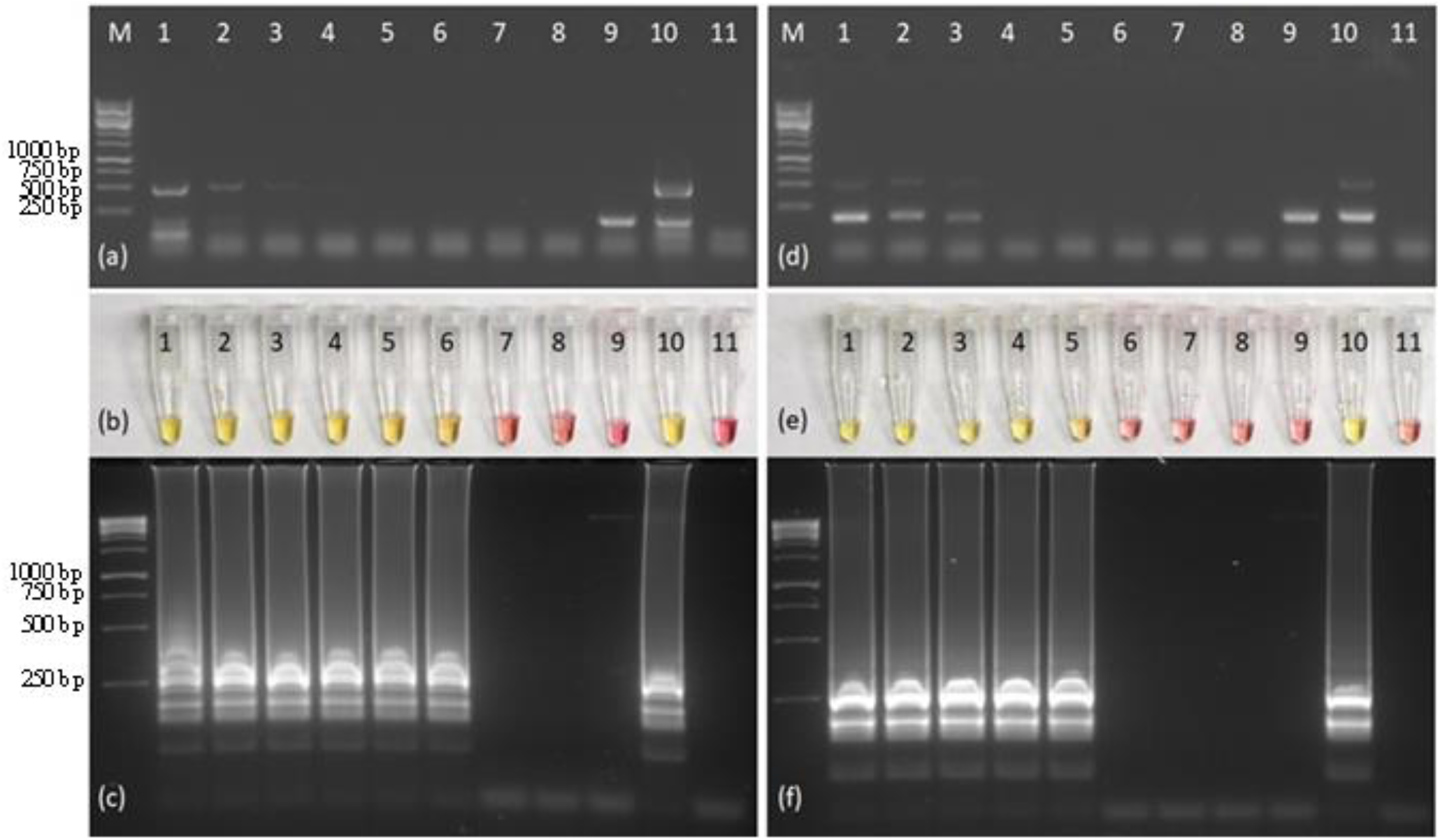
Relative sensitivity of RT-PCR and RT-LAMP for detecting GRSV and TSWV. Visualization of RT-PCR products amplified with specific primers for GRSV (a) and TSWV (d). RT-LAMP reactions performed using the P-73 primer set for GRSV (b) and the P-140 primer set for TSWV (e). Visualization of RT-LAMP products amplified with primer sets for GRSV (c) and TSWV (f). Lane M, 1 Kb DNA Ladder (Thermo Scientific); lane/tube 1, 500 ng/µl of total RNA extracted from virus-infected tissue; lanes/tubes 2 to 8, 10-fold serial dilutions of total RNA; lane/tube 9, negative control (healthy peanut); lane/tube 10, positive control; lane/tube 11, no template control.

To assess the field applicability of the developed RT-LAMP, samples of symptomatic orthotospovirus-infected tomato (n=1), peppers (n=4), and peanuts (n=9) from Florida and Georgia (USA), as well as peanuts (n=32) from Córdoba Province (Argentina), were tested for GRSV and TSWV using the designed primer sets.

The tomato sample tested negative for GRSV and TSWV. In contrast, pepper and peanut samples from the USA tested positive for TSWV, while peanut samples from Argentina tested positive for GRSV. Additionally, the tomato sample, which did not react with any of the primer sets, and the GRSV-positive peanut samples were tested for TCSV using a virus-specific primer set targeting the NSm protein gene on RNA M (Yilmaz et al., 2023). The symptomatic tomato sample tested positive for TCSV, while the peanut samples tested negative, ruling out mixed infections or reassortant isolates of GRSV (L_G_M_T_S_G_) in the analyzed peanut samples. These findings align with the viral species distribution reported in the respective regions (Adkins et al., 2015; de Breuil et al., 2021; Lai et al., 2021; Funderburk et al., 2022) and underscore the specificity of the RT-LAMP method.

This study introduces a rapid, species-specific, and highly sensitive RT-LAMP assay as a visual diagnostic tool for detecting and identifying GRSV and TSWV in important crops such as tomato, pepper, and peanuts. The P-73 primer set for GRSV and the P-140 set for TSWV target fragments of RNA S, enabling their use in combination with other primer sets to detect mixed infections or interspecific recombinant isolates. The high sensitivity of RT-LAMP renders it an ideal method for working with tissues or plants that yield low quantities of RNA during extraction, establishing it as an effective method for the early detection of plant viruses.

## Acknowledgements

We thank C. Vanderspool, J. Smith and R. Lewis for their assistance with inoculum maintenance, and to the growers who generously provided field samples.

## Compliance with ethical standards

## Ethical approval

This article does not contain any studies with human participants or animals performed by any of the authors.

